# Phosphorus Availability Modulates Flowering Time Through Subcellular Reprogramming of bGLU25 and GRP7 in Flowering Plants

**DOI:** 10.1101/2025.01.02.631137

**Authors:** Huikyong Cho, Ilyong Choi, Nadia Bouain, Amjad Nawaz, Luqing Zheng, Zaigham Shahzad, Federica Brandizzi, Seung Y Rhee, Hatem Rouached

## Abstract

The transition from vegetative to reproductive growth is vital for plant fitness and crop yield and is strongly influenced by nutrient availability. While nitrogen deficiency accelerates flowering, phosphorus (P) limitation delays it. However, the molecular basis for how P availability regulates flowering time remains unclear. Here, through genome-wide association mapping in Arabidopsis, we uncover genetic variation in *β-GLUCOSIDASE 25* (*bGLU25*) that modulates flowering under P-limited conditions. In P-sufficient environments, bGLU25 localizes to the endoplasmic reticulum. Under P limitation, however, bGLU25 translocates to the cytosol, a process mediated by P-regulated SERINE CARBOXY PEPTIDASE 50 (SCP50). In the cytosol, bGLU25 binds to JACALIN-LECTIN LIKE1 (AtJAC1), preventing the nuclear translocation of the *Flowering Locus C* (*FLC*) regulator GLYCINE-RICH RNA-BINDING PROTEIN 7 (GRP7). This cytosolic sequestration of GRP7 under P-deprivation elevates *FLC* expression, delaying flowering. Moreover, in the monocot rice, the homologs of bGLU25 also modulate flowering responses to P availability, indicating a conserved role for bGLU25 across flowering plants. Our findings provide a molecular framework for breeding strategies aimed at optimizing flowering time in response to P levels.

---

Regulating flowering time enables plants to align reproductive processes with favorable environmental conditions^1^. The transition to flowering reallocates resources from vegetative growth to reproduction, underscoring its critical role in overall plant success^2^. Beyond being essential for growth, nutrients can also function as signals that influence the timing of flowering^3–5^. To fine-tune their flowering time, plants evolved intricate molecular mechanisms involving the integration of multiple signaling pathways, including photoperiod, vernalization, gibberellin, autonomous, and age-related cues^6–9^. Among these, the *FLOWERING LOCUS C* (*FLC*) plays a pivotal role in regulating both autonomous and non-autonomous pathways^10,11^, helping plants optimize reproductive strategies in response to environmental stimuli, such as temperature and nitrous oxide levels^12^. Soil-based signals, including water and nutrient availability, also affect flowering time^4,5,13,14^. For example, P limitation typically delays flowering in annual plants^15,16^. Genetic studies revealed that mutations in the *PHOSPHATE TRANSPORTER TRAFFIC FACILITATOR1* (*PHF1*) gene^17^ lead to delayed flowering under low P conditions^18^, while mutants in *NITROGEN LIMITATION ADAPTATION* (*NLA*), which accumulate excess P, flower earlier^18,19^. Likewise, inactivation of *PHOSPHATE 1* (*PHO1*), a key regulator of P translocation from roots to shoots^20^, causes delayed flowering by repressing floral activators such as *FLOWERING LOCUS T* (*FT*)^4^. Notably, this delayed flowering phenotype in the *pho1* mutant can be reversed by P supplementation, indicating that P availability is a key determinant in the timing of flowering transition^4^. However, the molecular mechanisms by which plants sense and respond to P availability to regulate flowering time remain largely unknown. Understanding these processes could offer valuable insights into how plants adapt to nutrient-limited environments and could pave the way for improving crop yields in fluctuating nutrient conditions.

## bGLU25 is necessary for adjusting flowering time to P limitation in *Arabidopsis thaliana*

P limitation significantly delays flowering time (Figures 1 A and B), yet the genetic basis of this response remains unclear. To explore the genetic basis of this response, we investigated natural variation in flowering time under P limitation (-P) across 233 Arabidopsis accessions^21^ (Figure 1C, Table S1). This study uncovered a large variation in flowering time under -P (Figure 1C), with a broad-sense heritability (*H^2^*)^22^ of 0.41. Genome-wide association (GWA) mapping using an accelerated mixed model^23^ identified a significant association between single nucleotide polymorphisms (lead SNP = Chr3:947345; *P* = 8.18) with flowering time under -P (Figure 1D). To identify the gene responsible for this quantitative trait locus (QTL), we assessed the flowering time of Arabidopsis Col-0 (WT), two independent T-DNA insertion knock-out lines, and two lines overexpressing each of the five genes (*AT3G03620*, *AT3G03630*, *AT3G03640*, *AT3G03650*, and *AT3G03660*) located within 20 Kb of the GWA SNP, Chr3:947345 (Figure S1A). While knocking out or overexpressing *AT3G03620*, *AT3G03630*, *AT3G03650*, and *AT3G03660* genes did not result in significant changes in flowering time under -P compared to WT, two T-DNA knock-out lines for *AT3G03640* (*β-GLUCOSIDASE 25*, *bGLU25*), *bglu25-1* and *bglu25-2* (Figure S1B), flowered earlier (Figures 1E-H, Figure S1B). In contrast, overexpressing *bGLU25* (bGLU25-OE) delayed flowering under -P more than WT (Figures 1G-H, Figure S1C). Complementation of the early flowering phenotype in *bglu25-1* through *bGLU25* expression confirmed that *bGLU25* was responsible for the flowering time phenotype (Figures 1G-H, Figure S1C). At the molecular level, P limitation increased *bGLU25* expression by more than 2-fold in WT (Figure 1I). To examine the impact of -P on bGLU25 protein levels, we expressed *GFP::bGLU25* fusion protein driven by its native promoter in WT. Consistent with the transcriptional response, -P resulted in a significant increase in bGLU25 protein levels (Figure 1J). These results demonstrate that P limitation induces bGLU25 and bGLU25 plays a critical role in delaying flowering time in response to P limitation.

**Figure 1.**
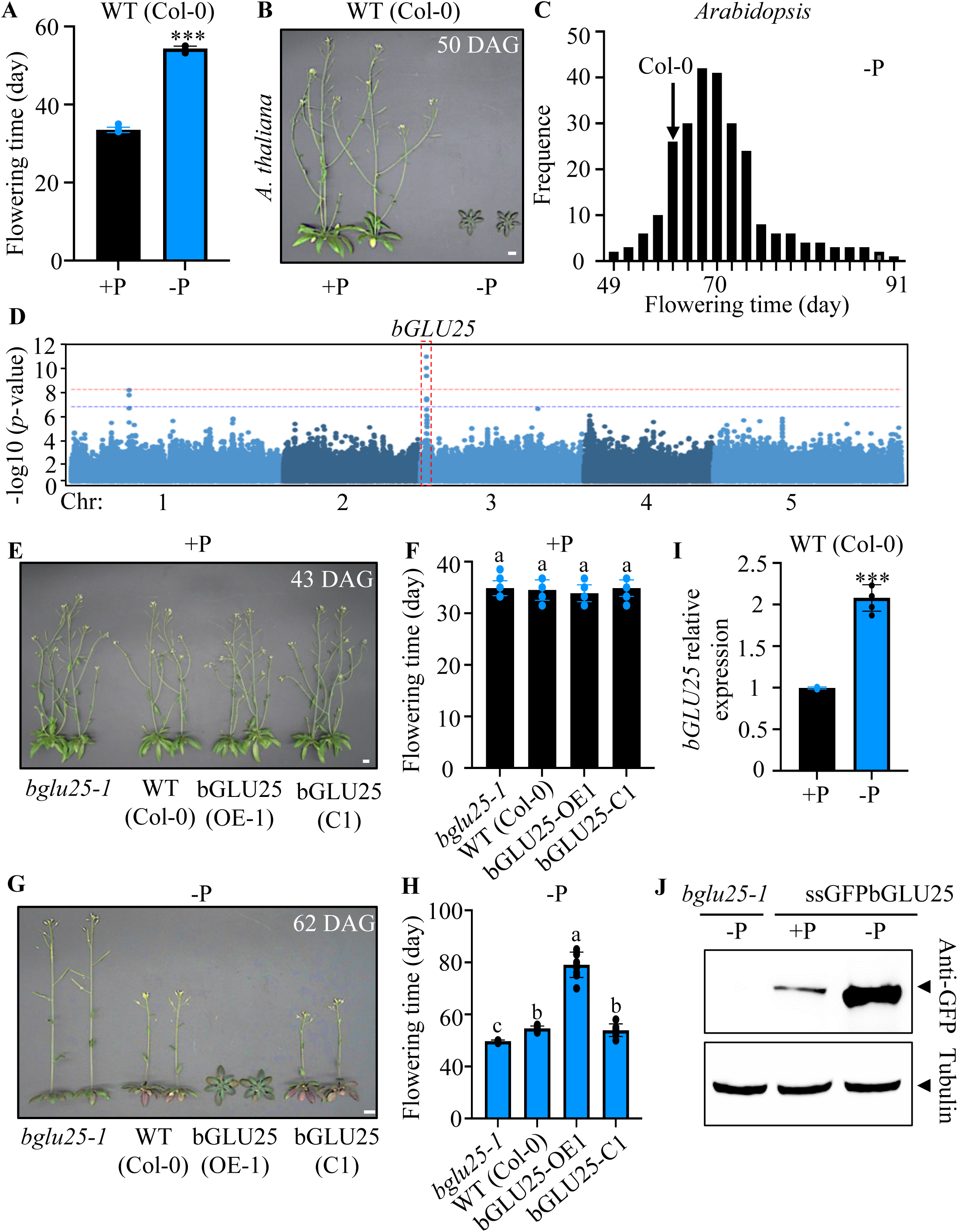
Genome-Wide Association (GWA) Analysis of *Arabidopsis* Flowering Time Under Phosphorus Limitation. **(A)** Flowering time of *Arabidopsis thaliana* (Col-0 accession) grown under phosphorus sufficient (+P) and deficient (-P) conditions. 18 plants from 3 independent experiments were analyzed. Student’s t test, \*\*\**p* < 0.001. **(B)** Representative images of wild-type (WT) plants (Col-0) grown under +P and -P conditions were taken 50 days after planting. (Scale bar: 1 cm) **(C)** Frequency distribution of mean of flowering time of 264 *A. thaliana* accessions grown in -P. **(D)** Manhattan plots from the GWA analysis show flowering time associations across accessions, with each dot representing the -log10(*p*) score of SNPs. The five *Arabidopsis* chromosomes are indicated. Red and dark dashed lines indicate the 0.05 False Discovery Rate and Bonferroni significance thresholds, respectively, with the *bGLU25* locus (AT3G03640) marked in red rectangle. **(E)** and (**G**) Representative images of WT, *bglu25-1*, complemented *bglu25-1* (bGLU25-C1), and 35S promoter-driven *bGLU25* overexpressor (bGLU25-OE1) plants grown under +P (**E**) and -P (**G**) conditions for 43 days (+P) or 62 days (-P) after germination. **(F)** and (**H**) Flowering time data for the plants shown in (**E**) and (**G**), with letters indicating significant differences at p < 0.05 (one-way ANOVA with Duncan post hoc test). **(I)** Relative mRNA abundance of *bGLU25* in 35-day-old Col-0 shoots was quantified by qRT-PCR, normalized to *Ubiquitin10*. Means (±95% confidence interval) of 18 plants from 3 independent experiments are shown. Student’s t test, \*\*\**p* < 0.001. **(J)** Immunoblotting of the bGLU25 protein in shoots of plants expressing ssGFPbGLU25, grown under +P and -P conditions for 35 days. Tubulin served as a loading control.

## Cytosolic localization of bGLU25 is linked to a delayed transition to flowering

To investigate how bGLU25 delays flowering time under P limitation, we first examined its subcellular localization. bGLU25 belongs to a β-glucosidase subfamily with eight members in *Arabidopsis*, all of which contain a signal peptide for endoplasmic reticulum (ER) targeting, suggesting ER localization^24^ (Figure S1D). To determine the subcellular localization of bGLU25, we expressed ssGFP::bGLU25 fusion protein, with GFP fused between the signal peptide and the remainder of bGLU25, transiently in *Nicotiana benthamiana* and stably in *Arabidopsis*. The fusion protein was functional, as it rescued *bglu25-1*’s early flowering phenotype under -P (Figures 1G-H), allowing us to assess bGLU25 localization across P levels. Under P sufficient conditions (+P), ssGFP::bGLU25 localized to the ER based on live imaging in *N. benthamiana* (Figure 2A) and density gradient centrifugation in *Arabidopsis* (Figure 2C). Surprisingly, under P deficient conditions (-P), ssGFP::bGLU25 predominantly localized to the cytosol in *N. benthamiana* leaves (Figure 2B). This observation was further corroborated in Arabidopsis cell fractionation assays (Figure 2C), demonstrating that bGLU25 is predominantly localized in the soluble fraction (cytosol) under -P, whereas it predominantly localizes to the ER under +P (Figure 2C). Next, we investigated whether the cytosolic translocation of bGLU25 affects flowering time. To test this, we generated a truncated version of bGLU25 (ΔbGLU25), lacking the 15 amino acids that contain the ER retention motif in the C-terminus (Figure S1D), and fused GFP between the signal peptide and the truncated protein. Confocal microscopy in *N. benthamiana* revealed that ΔbGLU25 was distributed in the cytosol (Figures 2D-E), which was confirmed in Arabidopsis using cell fractionation assays (Figure 2F). Furthermore, *Arabidopsis bglu25*-*1* mutant plants expressing ΔbGLU25 delayed flowering regardless of P availability (Figures 2G-H, Figures S3D-E), supporting the conclusion that the translocation of bGLU25 from the ER to the cytosol under P limitation contributes to delaying flowering in *Arabidopsis*.

**Figure 2.**
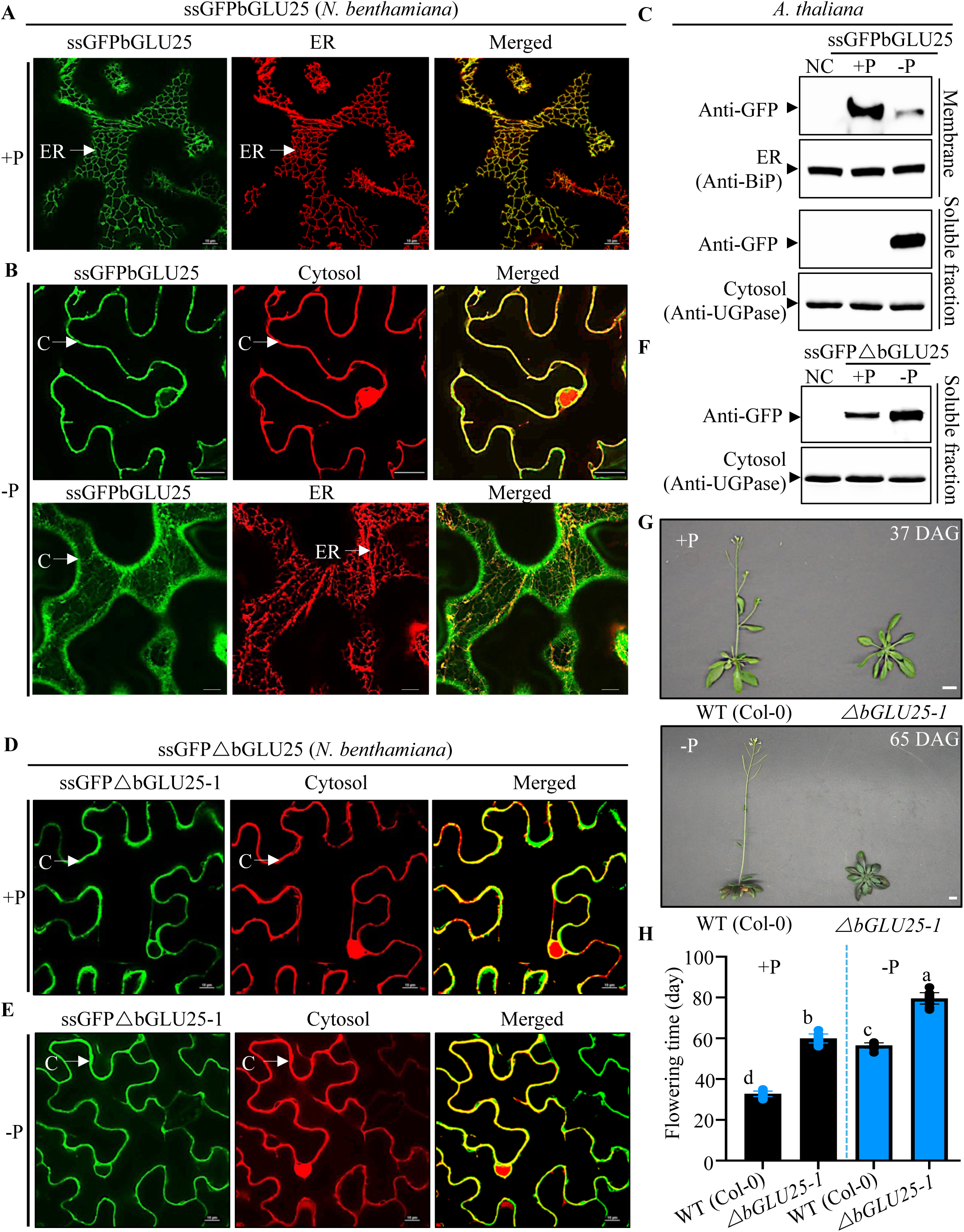
bGLU25 Relocates from the Endoplasmic Reticulum (ER) to the Cytosol Under Phosphorus Limitation. **(A)** Confocal microscopy images of *N. benthamiana* leaves from plants grown under +P conditions show the ssGFPbGLU25 signal of GFP (green fluorescence, left), the ER marker HDEL-RFP (red fluorescence, middle), and an overlay of both images (right). Scale bar: 10 μm. ER, Endoplasmic Reticulum. **(B)** Confocal microscopy images of *N. benthamiana* leaves from plants grown under -P conditions show the ssGFPbGLU25 signal of GFP (green fluorescence, left), the cytosol marker (red fluorescence, middle) or ER marker HDEL-RFP (red fluorescence, middle), and an overlay of the images (right). Scale bar: 10 μm. C, cytosol. ER, Endoplasmic Reticulum. **(C)** Immunoblotting of the ssGFPbGLU25 protein in the cytosol and ER-enriched fraction of plants grown under +P and -P conditions. BiP (luminal binding protein, Binding Immunoglobulin Protein) and UGPase (UDP-glucose pyrophosphorylase) were used as markers, while non-transformed plants served as negative controls (NC). **(D)** and (**E**) Confocal microscopy images of *N. benthamiana* leaves from plants grown under +P (**D**) and -P (**E**) conditions show the ssGFPΔbGLU25 signal of GFP (green fluorescence, left), the cytosol marker (red fluorescence, middle), and an overlay of both images (right). Scale bar: 10 μm. C, cytosol. **(E)** Immunoblotting of the ssGFPΔbGLU25 protein in ssGFPΔbGLU25 transgenic lines, with UGPase as a marker for the soluble fractions, with NC as negative controls. **(F)** Images of wild-type and plants grown under +P or -P conditions for 37 days (+P) or 65 days (-P). Scale bar: 1 cm. **(G)** Flowering time data for the plants shown in (**G**). 18 plants from 3 independent experiments were analyzed. Letters indicate significant differences at p < 0.05 (one-way ANOVA with Duncan post hoc test).

## Release of bGLU25 from the ER to cytosol requires the SCP50 protein peptidase

To investigate the mechanism driving bGLU25 translocation to the cytosol under P limitation, we used a candidate gene approach, focusing on P-responsive carboxypeptidases documented in the literature^25^. These included *LEUCINE AMINOPEPTIDASE 1* (AT2G24200), *LEUCINE AMINOPEPTIDASE 2* (AT4G30920), *SERINE CARBOXYPEPTIDASE 20* (AT4G12910), and *SERINE CARBOXYPEPTIDASE 50* (*SCP50*, AT1G15000) ^25^. We examined the flowering time of knockout mutants of these genes under varying P conditions. Only the genetic inactivation of *SCP50* (*scp50-1*) resulted in early flowering, similar to the phenotype observed in *bglu25* knock-out plants (Figures 3A-B, Figures S2A-B). Quantitative real-time PCR analysis showed that the transcript levels of *SCP50* are induced by P limitation (Figure 3C). To investigate how SCP50 protein levels and subcellular localization are regulated by P limitation, we generated SCP50-GFP fusion constructs. Under P-limited conditions, SCP50-GFP protein levels were significantly increased (Figure 3D), and the protein localized to the ER (Figure 3E). To test whether SCP50 affects the trafficking of bGLU25, we expressed ssGFP::bGLU25 in *scp50-1* mutant plants. In these mutants, cytosolic localization of bGLU25 under - P was abolished, with bGLU25 confined to the ER (Figures 3F-G). These results support the notion that SCP50 mediates the translocation of bGLU25 from the ER to the cytosol in response to P limitation, potentially delaying flowering.

**Figure 3.**
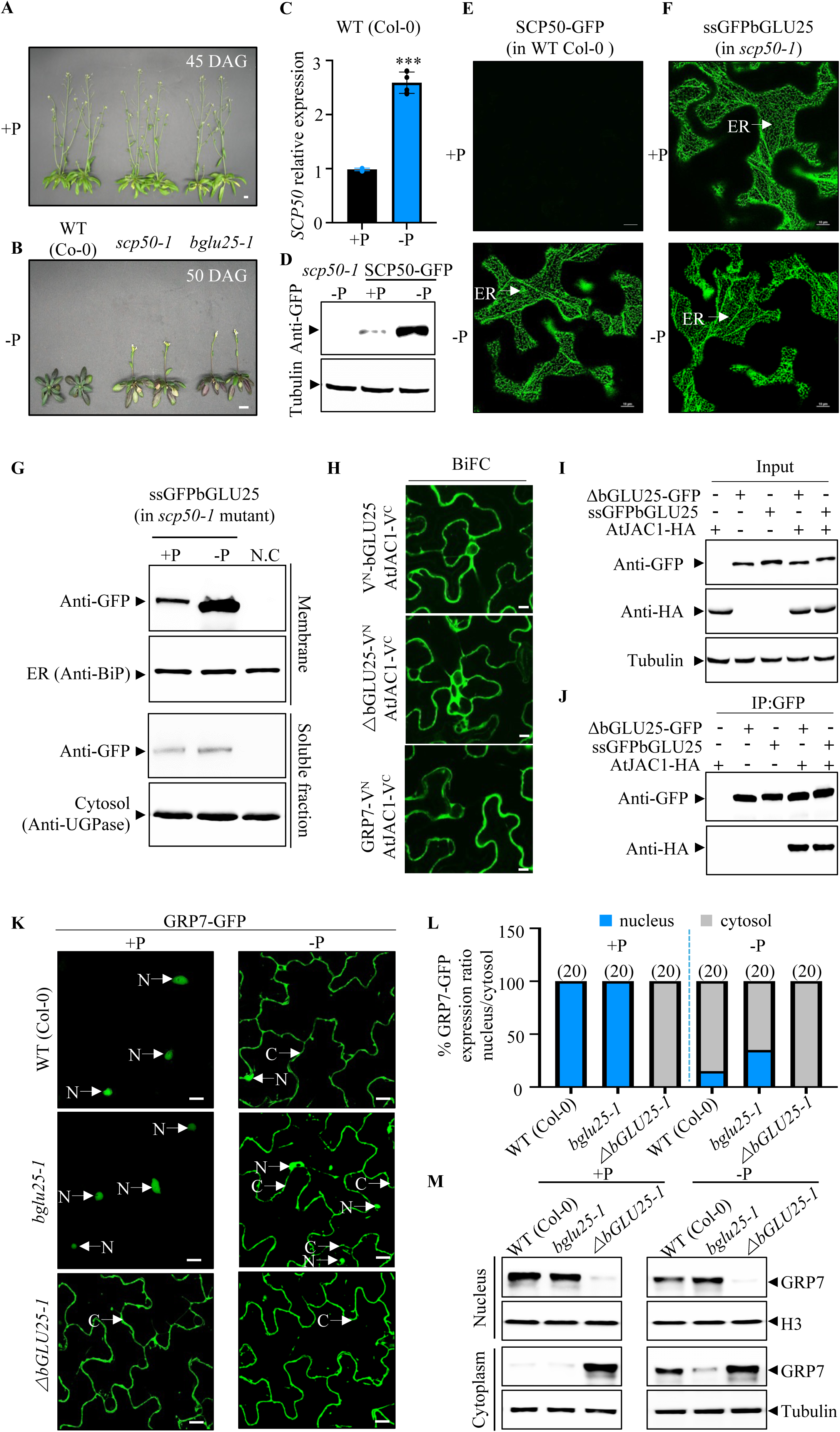
SCP50 Facilitates bGLU25 Cytosolic Translocation Under Phosphorus Limitation, Regulating GRP7 Localization via AtJAC1 Interaction. **(A)** and (**B**) Representative images of WT (Col-0), *scp50* mutant (*scp50-1*), and *bglu25* mutant (*bglu25-1*) plants grown under phosphorus-sufficient (+P) and phosphorus-deficient (-P) conditions for 45 and 50 days, respectively (scale bar = 1 cm). **(B)** Relative mRNA abundance of *SCP50* in shoots of 35-day-old *Arabidopsis* (Col-0) plants grown under +P and -P conditions, quantified by qRT-PCR and normalized to *Ubiquitin10*. Means ±95% confidence interval are shown. Individual measurements were obtained from the analysis of shoots collected from a pool of 6 plants. Student’s t test, \*\*\**p* < 0.001. **(C)** Western blot showing SCP50 protein levels in *A. thaliana* shoots of plants grown under +P and -P conditions, with GFP-tagged SCP50 protein detected using anti-GFP antibodies. *scp50-1* mutant lines served as specificity controls. Tubulin was used as a loading control. **(D)** and (**F**) Confocal microscopy images of *A. thaliana* leaves displaying the SCP50-GFP signal in a wild-type background (**E**) and the ssGFPbGLU25 signal in the *scp50-1* mutant background (**F**), grown under +P and -P conditions. Scale bar: 10 μm. ER, Endoplasmic Reticulum. **(G)** Immunoblotting of the ssGFPbGLU25 protein in the soluble (cytosol) and membrane (ER-enriched) fractions of plants grown under +P and -P conditions. BiP was used to detect ER markers, and UGPase was used for cytosol markers. Non-transformed plants served as controls (NC). **(H)** BiFC assays showing the interaction between bGLU25 or ΔbGLU25 and AtJAC1 transiently expressed in *N. benthamiana*. AtJAC1 and GRP7 were used as positive controls. **(I)** and (**J**) Co-IP assays for the interactions between bGLU25 or ΔbGLU25 and AtJAC1. **(K)** Confocal microscopy images from *A. thaliana* showing GRP7-GFP signal in nucleus and cytosol of leaves of wild-type, *bglu25-1* mutant, and *ΔbGLU25-1* lines grown under +P and -P conditions (Scale bar: 10 μm). N, Nucleus. C, Cytosol. **(L)** Quantification of GRP7-GFP distribution between the nucleus and cytosol, as shown in (**K**). Fluorescence intensities in leaf cells from WT, *bglu25-1*, and *ΔbGLU25-1* are presented as percentages based on the average of 20 measurements. **(M)** Western blot analysis confirms GRP7-GFP localization across different lines under +P and -P conditions (**K**).

## bGLU25 is an inactive enzyme that regulates flowering time

Arabidopsis bGLU25 belongs to a large family of β-glucosidase enzymes, which typically hydrolyze terminal, non-reducing β-D-glucosyl residues from glyco-conjugates to produce glucose^24^. Interestingly, bGLU25 lacks key catalytic residues and motifs required for enzyme activity compared to active β-glucosidases such as bGLU15, bGLU22, and bGLU23 (Figure S1D). Based on this observation, we hypothesized that bGLU25 is an inactive enzyme with a distinct molecular function. To test this, we assayed β-glucosidase activity of bGLU25 using several substrates (esculin, ONPG, isoquercetin, astragalin). Unlike bGLU23, a confirmed active β-glucosidase (Figure S2C), bGLU25 showed no catalytic activity with any of these substrates, supporting the hypothesis that bGLU25 is an inactive β-glucosidase. We then asked whether bGLU25 activity could be restored by substituting a key active site residue. We generated a bGLU25^G420E^ mutant by replacing glycine at position 420 with glutamic acid, a residue critical for enzyme activity for β-glucosidases^24^. This mutant form displayed modest β-glucosidase activity, producing detectable amounts of product from various substrates such as esculin, ONPG, isoquercetin, and astragalin (Figure S2C). To assess whether reviving the β-glucosidase activity of bGLU25 influences flowering time, we expressed both the inactive WT bGLU25 and the partially active bGLU25^G420E^ mutant, driven by the native promoter in the *bglu25* mutant background. *bglu25* plants expressing either bGLU25 or bGLU25 ^G420E^ exhibited similar flowering time (Figures S2D-F), indicating that reviving β-glucosidase activity of bGLU25 does not interfere with its function in regulating flowering time. These findings suggest that inactive enzymes, like bGLU25, can serve critical regulatory functions in plant development, independent of their catalytic activity.

## bGLU25 interacts with AtJAC1 to regulate GRP7 localization and flowering time in *Arabidopsis* via P-dependent mechanisms

To gain insights into how the cytosolic translocation of bGLU25 under -P delays flowering, we searched for interacting proteins of bGLU25. Using both the full-length bGLU25 and ΔbGLU25 variants as bait proteins in a yeast two hybrid screen, we identified the same set of interacting partners for both baits (Table S2), indicating that the C-terminus is not responsible for interaction specificity. Among the interacting partners, JACALIN-LECTIN LIKE1 (AtJAC1), a known repressor of flowering in Arabidopsis, emerged as a high-confidence interactor of bGLU25 (Figure S3A). Loss of function mutations in *AtJAC1* cause early flowering, while its overexpression delays flowering^26^. AtJAC1 modulates flowering time by controlling the nucleocytoplasmic distribution of GLYCINE-RICH RNA-BINDING PROTEIN7 (GRP7)^26^. GRP7 regulates the expression of *Flowering Locus C* (*FLC*), a central repressor of flowering, through processing of *FLC* antisense transcripts^26^. The interaction between bGLU25 and AtJAC1 was further validated through bimolecular fluorescence complementation (BiFC) assays in *N. benthamiana* (Figure 3H, Figure S3B) and co-immunoprecipitation experiments (Figures 3I-J), confirming a direct protein-protein interaction. To test whether the interaction between bGLU25 and AtJAC1 affects the nucleocytoplasmic distribution of GRP7 in a P-dependent manner, we expressed GRP7-GFP in *N. benthamiana* and in various Arabidopsis backgrounds (Col-0, *bglu25-1*, and *ΔbGLU25-1*). In both, *N. benthamiana* (Figure S3C), and in wild-type Arabidopsis, GRP7 predominantly localizes to the nucleus under +P conditions but shifts predominantly to the cytosol under -P conditions (Figures 3K-L). A similar distribution of GRP7 was observed in *bglu25-1* mutants under both +P and -P conditions, though propensity of nuclear localization of GRP7 under -P was slightly higher in *bglu25-1* mutants compared to wild-type plants (Figures 3K-L). Remarkably, in *ΔbGLU25-1* plants, GRP7 remained predominantly cytosolic regardless of P availability (Figures 3K-L). In *ΔbGLU25-1* plants, GRP7 remained cytosolic under both +P and -P conditions, which was confirmed by cell fractionation assays (Figure 3M). These results highlight the key role of bGLU25 in regulating GRP7 localization in response to P availability.

To confirm that the bGLU25-AtJAC1-GRP7 complex operates within the same pathway regulating flowering time in response to P availability, we tested for genetic interactions among bGLU25, AtJAC1, and GRP7. Under P limitation, *bglu25-1, atjac1-1,* and *atjac1-1;bglu25-1* single and double mutants all flowered early (Figures 4A-B). However, *ΔbGLU25* plants, which keep bGLU25 in the cytosol and delays flowering, was epistatic to *atjac1-1,* indicating that bGLU25 and AtJAC1 function in a shared genetic pathway (Figures 4A-B, Figures S3D-E). Moreover, the late-flowering phenotype of *grp7-1* mutants persisted regardless of P availability (Figures 4C-D, Figures S3D-E) and was epistatic to both *bglu25-1* and *atjac1-1,* indicating that GRP7 acts downstream of bGLU25 and AtJAC1. Consistently, the *grp7-1;bglu25-1;atjac1-1* triple mutant also flowered late. These findings suggest that bGLU25 acts upstream in this regulatory pathway involving AtJAC1 and GRP7. Gene expression analysis showed that *bglu25-1* and *atjac1-1* mutants repress *FLC* levels, whereas *grp7-1* single mutants and the *grp7-1;bglu25-1;atjac1-1* triple mutants exhibited high levels of *FLC* mRNA, consistent with their respective flowering phenotypes (Figure 4E). To further confirm that bGLU25 influences flowering time through *FLC* expression modulation (Figure 4E), we crossed the late-flowering *ΔbGLU25* mutant with the early-flowering *flc-1* mutant. The resulting *ΔbGLU25;flc-1* lines produced early-flowering plants, indicating that FLC acts downstream of bGLU25 and that ΔbGLU25’s role on flowering time requires FLC (Figures 4F-G, Figure S3D). Together, these findings emphasize the crucial role of bGLU25, AtJAC1, and GRP7 in regulating flowering time through FLC in a P-dependent manner.

**Figure 4.**
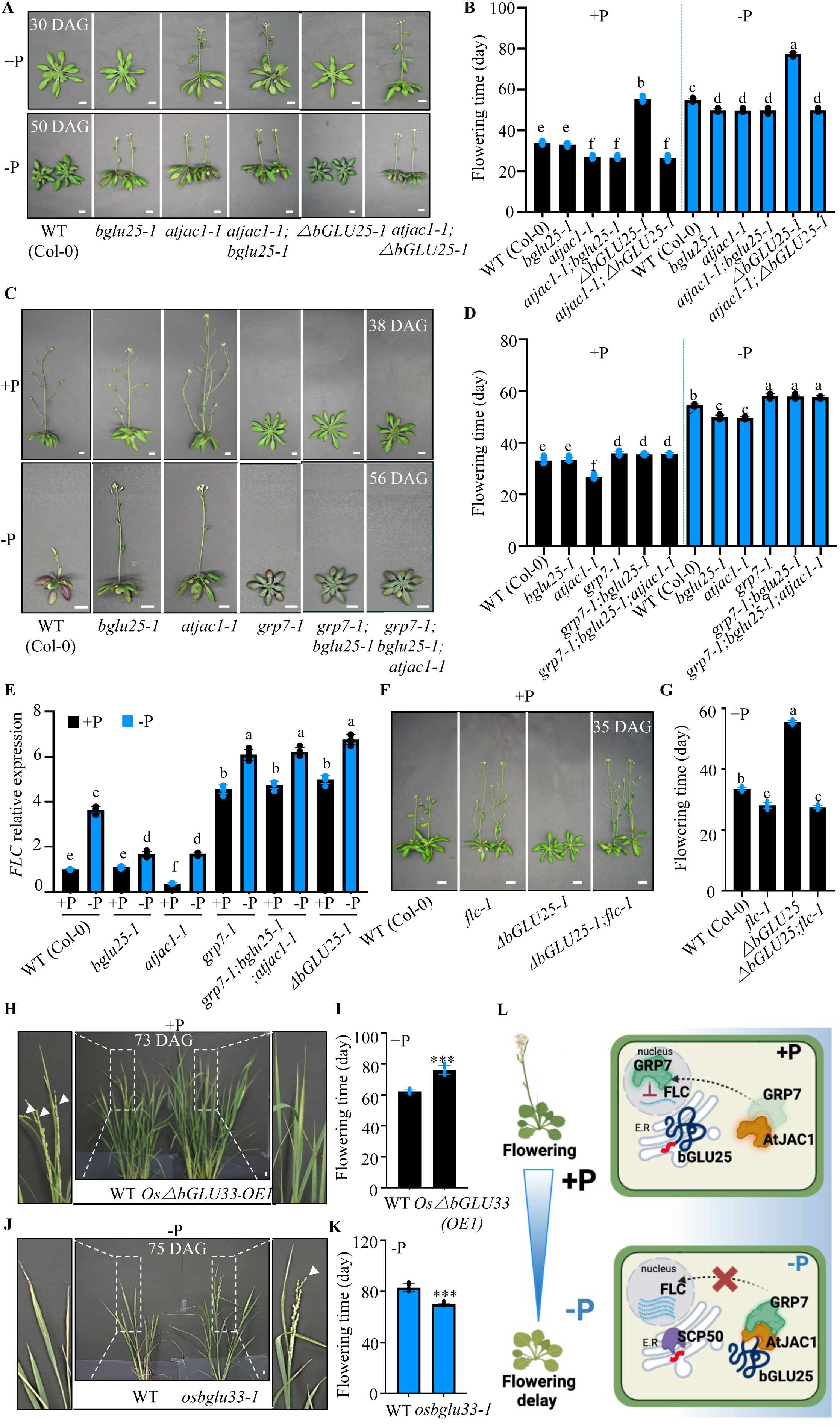
bGLU25’s Function in Flowering Time Regulation Is Conserved in Rice. **(A)** Representative images of wild-type (WT), *bglu25-1*, *atjac1-1*, *ΔbGLU25-1*, the double mutants *atjac1-1;bglu25-1*, and *atjac1-1;ΔbGLU25-1* grown under +P and -P conditions, taken at 30 days (+P) and 50 days (-P). (Scale bar: 1 cm). **(B)** Flowering time data for the plants shown in (**A**), with letters indicating significant differences at p < 0.05 (one-way ANOVA with Duncan post hoc test). **(C)** Representative images of WT, *bglu25-1*, *atjac1-1*, *grp7-1*, *grp7-1;bglu25-1*, and *grp7-1;bglu25-1;atjac1-1*plants, grown under +P and -P conditions for 38 days (+P) or 56 days (-P). (Scale bar: 1 cm). **(D)** Flowering time data for the plants shown in (**C**), with letters indicating significant differences at p < 0.05 (one-way ANOVA with Duncan post hoc test). **(E)** Relative mRNA abundance of *FLC* in the shoots of the lines mentioned in (**C**) and *ΔbGLU25-1* plants was quantified by qRT-PCR, normalized to *Ubiquitin10*. Data are presented as means (±95% confidence interval). Individual measurements were obtained from the analysis of shoots collected from a pool of six 35-day-old plants. Statistical analysis was performed using Student’s t-test, ***p < 0.001. **(F)** Representative images of *A. thaliana* wild-type (WT), *flc-1*, *ΔbGLU25-1*, and the double mutant *ΔbGLU25-1; flc-1.* (Scale bar: 1 cm). **(G)** Flowering time data for the plants shown in (**F**), 18 plants from 3 independent experiments were analyzed, with letters indicating significant differences at p < 0.05 (one-way ANOVA with Duncan post hoc test). **(H)** and (**J**) Images of *Oryza sativa* (rice) wild-type (WT) and *ΔbGLU33* overexpressor plants grown under +P conditions (**H**), as well as WT and *osbglu33-1* mutant plants grown under -P conditions (**J**). (Scale bar: 1 cm). White arrows indicate the rice flowers. **(I)** and (**K**) Flowering time data for the lines shown in (**H**) and (**J**), respectively. 18 plants from 3 independent experiments were analyzed, with letters indicating significant differences at p < 0.05 (one-way ANOVA with Duncan post hoc test). (**L**) Schematic model illustrating the role of the bGLU25-AtJAC1-GRP7 axis in regulating flowering time through the *FLC* signaling pathway in a P-dependent manner.

## bGLU25-dependent regulation of flowering time is likely conserved in rice

The critical role of bGLU25 in regulating flowering time in response to P availability prompted us to investigate its conservation across plants. The bGLU gene family is widely represented in the genomes of most plant taxa, including grasses such as rice, indicating a potential evolutionary significance in flowering time modulation and potential for translation of this mechanism into crops. We hypothesized that analogous alterations in bGLU25 homologs in rice would similarly affect flowering time in a P-dependent manner. We identified OsbGLU33 (LOC_Os09g33710) as a close homolog of *Arabidopsis* bGLU25, exhibiting 52% protein similarity and 38% identity^27^ (Figure S4A). Remarkably, *OsbGLU33*, like *bGLU25*, is induced by P limitation (Figure S4B). To investigate its function, we created independent lines with CRISPR-Cas9-mediated inactivation of *OsbGLU33* and lines expressing the cytosolic form *OsΔbGLU33*, achieved by removing 10 amino acids at the C-terminus that include the putative ER retention motif (Figure S4A). Like *Arabidopsis*, under +P conditions, the full length OsbGLU33 localizes to the ER, while under -P conditions, it is found in the cytosol (Figure S4C). The truncated OsΔbGLU33 is consistently cytosolic regardless of P availability (Figure S4D). We then compared the flowering time of wild-type rice, *osbglu33* mutants, and *OsΔbGLU33* lines under varying P regimes. Notably, OsΔbGLU33 overexpressor lines exhibited delayed flowering, even under sufficient P conditions (Figures 4H-I, Figures S4E-F), while *osbglu33-1* mutants flowered earlier, mirroring the effects observed in *Arabidopsis* (Figures 4J-K, Figures S4E-F). These results indicate genetic conservation of the bGLU25-dependent regulatory pathway in rice, suggesting its pivotal role in modulating flowering time across diverse plant species.

## DISCUSSION

P is a finite resource^28^, and its scarcity poses a significant threat to global food security ^29^. P limitation is particularly detrimental to agriculture, as it delays the transition from vegetative to reproductive growth, which directly impacts crop yield^2^. Understanding how plants adapt to this limitation is crucial for developing strategies that improve crop resilience to nutrient-poor environments. Our study examining the genetic basis and natural variation of flowering time under P limitation in *Arabidopsis thaliana* reveals that delayed flowering under P limitation represents an adaptive response to optimize survival in nutrient-limited conditions. This finding underscores the importance of postponing flowering as a strategy to enhance plant fitness in P-deficient soils^16^. Through our identification of bGLU25 as a central regulator of flowering time in response to P availability (Figure 4L), we have uncovered key mechanisms underlying this adaptive process. Under sufficient P conditions, bGLU25 is localized to the ER, while the AtJAC1-GRP7 complex resides in the cytosol (Figure 4L). As the plant approaches flowering, GRP7 translocates to the nucleus to repress the floral repressor FLC, thereby promoting flowering. However, under P-deficient conditions, bGLU25 is trasnslocated to the cytosol via the P-inducible protein SCP50, stabilizing its interaction with the AtJAC1-GRP7 complex via AtJAC1 and limiting GRP7’s entry to the nucleus (Figure 4L). This retention of GRP7 in the cytosol maintains high *FLC* levels, thereby delaying flowering (Figure 4L). Notably, the bGLU25-dependent regulatory mechanism appears conserved in rice, suggesting that manipulating this pathway could significantly improve crop performance and ensure agricultural productivity in the face of global nutrient challenges, with the potential to contribute meaningfully to future food security.

## MATERIALS AND METHODS

### Plant materials and growth conditions

Seeds of *Arabidopsis thaliana* wild type (Columbia ecotype, Col-0, stock number CS60000) and various knock-out mutant lines for bGLU25 (AT3G03640), specifically *bglu25-1* (SALK_128790) and *bglu25-2* (SALK_128792), were sourced from The Arabidopsis Biological Resource Center (ABRC, USA). In addition to the *bGLU25* lines, knock-out mutant lines for *LEUCINE AMINOPEPTIDASE 1* (AT2G24200, SALK_086638), L*EUCINE AMINOPEPTIDASE 2* (AT4G30920, SALK_208947), *SERINE CARBOXYPEPTIDASE 20* (AT4G12910, SAIL_392_C07), and *SERINE CARBOXYPEPTIDASE 50* (SCP50, AT1G15000; *scp50-1*: SALK_092741 and *scp50-2*: SAIL_87_E12), were also obtained from ABRC. Moreover, knock-out lines for the genes *MULTIDRUG AND TOXIC COMPOUND EXTRUSION (MATE)* (AT3G03620, SALK_113658); *CYSTEINE SYNTHASE 26* (AT3G03630, SALK_205530); *EMBRYO SAC DEVELOPMENT ARREST 5* (AT3G03650, SAIL_389_C05); *WUSCHEL RELATED HOMEOBOX 11* (AT3G03660, SALK_004777); *AtJAC1* (AT3G16470; *atjac1-1*: SALK_000461 and *atjac1-2*: SALK_112556), GRP7 (AT2G21660; *grp7-1*: SALK_039556 and *grp7-2*: SALK_113110), and *FLC* (AT5G10140; *flc-1*: SALK_041126 and *flc-2*: SALK_072590) were included in this study, with their homozygosity verified by PCR. Double mutants involving *bglu25* crossed with *atjac1-1*, *grp7-1*, and *flc-1* were created from the respective single mutants. For the GWAS analysis, we utilized a selection of 233 *A. thaliana* accessions from the RegMap panel (ABRC accession number CS77400), which were stored at 4°C until needed, with their identifiers listed in Table S1. Plants were grown hydroponically under nonsterile conditions. Seeds were germinated directly on top of pierced Eppendorf tubes filled with sand and placed on rafts floating in a hydroponic solution. Plants were germinated and grown for 1 week in tap water then in the following nutrient solution: 0.5 mM KNO_­_; 1 mM MgSO_­_; 1 mM KH_­_PO_­_; 0.25 mM Ca(NO3)2; l00 μM NaFeEDTA; 30 μM H_­_BO_­_; l0 μM MnCl_­_; l μM CuCl_­_; 1 μM ZnCl_­_; 0.1 μM (NH_­_)6Mo_­_O_­_; and 50 μM KCl. For P limitation treatment, 1 mM KH_­_PO_­_was replaced by K_­_SO_­_. The nutrient solution was renewed every 4 d and on the day before the beginning of every treatment. Plants were grown in a growth chamber under the following environmental conditions: light/dark cycle of 16h/8h, light intensity of ∼120 μmol·m^−2^·s^−1^, temperature of 22/20°C, and RH of 75%. Flowering time was recorded when floral buds became visible in the center of the rosette, with data typically collected for six plants per accession and averaged for analysis from three experiments. For the rice experiments, we utilized a controlled-environment chamber with a 14-hour light and 10-hour dark cycle at a light intensity of 200 µmol photons·m²·s⁻¹, maintaining temperatures of 28 °C during the day and 25 °C at night, with 80% relative humidity. Briefly, approximately 50 rice seeds were cleaned, soaked in deionized water for 24 hours, and then germinated on moistened fabric for 48 hours. Seven-day-old seedlings, including wild-type, *osbglu33* mutants, and *Os.6.bGLU33* overexpression lines (*Os.6.bGLU33-OE1* and *Os.6.bGLU33-OE2*), were transplanted into a hydroponic growth system containing nutrient solutions. The solutions were adjusted to pH 5.9 and renewed every three days until the flowering time was recorded, as described in our recent publication^32^.

### Creation of bGLU25 and OsbGLU33 transgenic and knockout lines

To create Arabidopsis bGLU25 and ΔbGLU25 overexpression lines (*bGLU25-OE1*, *bGLU25-OE2*, *ΔbGLU25-1* and *ΔbGLU25-2*), the full-length coding sequence was amplified using PCR and cloned into the pDONR 223 vector. The LR clonase reaction transferred the *bGLU25* clone into the pMDC32 vector under the cauliflower mosaic virus 35S promoter, followed by overexpression in the *bglu25-1* mutant background. A G420 to E substitution in *bGLU25* was introduced via site-directed mutagenesis^30^ and expressed in the *bglu25-1* background, driven by the native promoter. For plants overexpressing *ssGFPbGLU25* or *ssGFPΔbGLU25*, the sequences were synthesized (Genscript) and cloned into the pUC57 vector. An LR clonase reaction transferred these into the pMDC32 vector, where the 35S promoter was replaced with the native *bGLU25* promoter using SbfI and ApaI restriction sites (see Table S3). These constructs were inserted into the binary vector pCAMBIA1301 and transformed into Agrobacterium tumefaciens strain GV3101 for the floral dip^31^ transformation of Arabidopsis (WT (Col-0), *bglu25-1* or *scp50-1* mutants). Transgenic plants were selected for hygromycin resistance, and only T3 homozygous progeny from heterozygous T1 plants, whose T2 progeny showed a 1:3 segregation ratio for antibiotic sensitivity, were analyzed. For rice, *OsbGLU33* knockout lines in *Oryza sativa* cv *Nipponbare* were generated using the CRISPR-Cas9 system^33^. Two guide RNAs targeting exons 1 and 2 of the *OsbGLU33* gene were selected from the CRISPR-PLANT database and inserted into the pRGEB32 vector^34^, producing two independent homozygous mutants, *osbglu33-1* and *osbglu33-2*, confirmed by Sanger sequencing for functional analysis. For overexpression lines of *OsbGLU33*, the full coding sequence or the version without the last 10 C-terminal amino acids was amplified and cloned into the pDONR223 vector. The LR clonase reaction transferred these into the pMDC32 vector under the cauliflower mosaic virus 35S promoter, creating the *35S-ssGFPOsbGLU33* and *35S-ssGFPOs.6.bGLU33* constructs. These were introduced into *Agrobacterium tumefaciens* and used to transform rice. Taq-Man PCR assessed copy numbers in the T0 generation, and single-copy plants were propagated at the MSU plant growth facility, with further studies on homozygous T3 plants.

### GWA mapping

GWAS was conducted using the GWA-portal (https://gwas.gmi.oeaw.ac.at/)^23^, employing a mixed model algorithm (AMM) that considers population structure^23^ along with SNP data from the RegMap panel^21,35,36^. To address multiple hypothesis testing, false discovery rate (FDR) was calculated using the Benjamini-Hochberg method^37^, with an FDR threshold of 5% set to identify significant associations.

### Yeast two-hybrid analysis

Yeast two-hybrid (Y2H) screening, and the direct 1-by-1 interaction assays confirmation, were performed by Hybrigenics Services, S.A.S., Evry, France (http://www.hybrigenics-services.com). Coding sequences of full-length and a C-terminal deletion mutant (aa 1-516) of *A. thaliana* bGLU25 (NM_111235.4) were PCR-amplified and cloned, in frame with the LexA DNA binding domain, into pB27 as a C-terminal fusion to LexA (LexA-bait fusion). The constructs were checked by sequencing the entire inserts and used as baits to screen a random-primed *A. thaliana* one week-old seedling cDNA library constructed into pP6. pB27 and pP6 derive from the original pBTM116 ^38^ and pGADGH ^39^ plasmids, respectively. For the screen with full-length bGLU25, 71 million clones (7-fold the complexity of the library) were screened using a mating approach with YHGX13 (Y187 ade2-101::loxP-kanMX-loxP, mata) and L40DGal4 (mata) yeast strains as previously described^40^. 160 His+ colonies were selected on a medium lacking tryptophan, leucine and histidine. For the C-terminal deletion mutant, 88 million clones (9-fold the complexity of the library) were screened using the same mating approach. 149 His+ colonies were selected on a medium lacking tryptophan, leucine and histidine. The prey fragments of the positive clones were amplified by PCR and sequenced at their 5’ and 3’ junctions. The resulting sequences were used to identify the corresponding interacting proteins in the GenBank database (NCBI) using a fully automated procedure. A confidence score (PBS, for Predicted Biological Score) was attributed to each interaction as previously described^41^.

The PBS relies on two different levels of analysis. First, a local score considers the redundancy and independency of prey fragments, as well as the distribution of reading frames and stop codons in overlapping fragments. Second, a global score considers the interactions found in all the screens performed at Hybrigenics using the same library. This global score represents the probability of an interaction being nonspecific. For practical use, the scores were divided into four categories, from A (highest confidence) to D (lowest confidence). A fifth category (E) specifically flags interactions involving highly connected prey domains previously found several times in screens performed on libraries derived from the same organism. Finally, several of these highly connected domains have been confirmed as false positives of the technique and are now tagged as F. The PBS scores have been shown to positively correlate with the biological significance of interactions^42,43^.

For the direct 1-by-1 interaction, the coding sequence for amino acids 38-193 of *A. thaliana* T1M15.190 (NM_122086.2) was PCR-amplified and cloned in frame with the Gal4 DNA binding domain (DBD) into plasmid pB66 as a C-terminal fusion to Gal4 (Gal4-bait fusion). pB66 derives from the original pAS2ΔΔ vector ^40^. The coding sequences of full-length and a C-terminal deletion mutant (aa 1-516) of *A. thaliana* bGLU25 (NM_111235.4) were PCR-amplified and cloned in frame with the LexA DNA binding domain (DBD) into pB27 as a C-terminal fusion to LexA (LexA-bait fusion). pB27 derives from the original pBTM116 vector^38^. Fragments of interacting proteins were extracted from the ULTImate Y2H™ screenings of T1M15.190 and bGLU25 with an Arabidopsis one week-old seedlings library. Prey fragments are cloned in frame with the Gal4 activation domain (AD) into plasmid pP6, derived from the original pGADGH^39^. The Gal4 bait construct was transformed in the yeast haploid cells CG1945 (mata) and the LexA bait construct was transformed in the yeast haploid cells L40deltaGal4 (mata), respectively. The prey constructs were transformed into YHGX13 (Y187 ade2-101::loxP-kanMX-loxP, mata). The diploid yeast cells were obtained using a mating protocol with both yeast strains^40^. The interaction assays are based on the HIS3 reporter gene (growth assay without histidine). As negative controls, the bait plasmids were tested in the presence of empty prey vector and all prey plasmids were tested with the empty bait vector. The interaction between SMAD (Suppressor of Mothers against Decapentaplegic) and SMURF (SMad Ubiquitination Regulatory Factors) is used as positive control^44^. Controls and interactions were tested in the form of streaks of three independent yeast clones on DO-2 and DO-3 selective media. The DO-2 selective medium lacking tryptophan and leucine was used as a growth control and to verify the presence of the bait and prey plasmids. The DO-3 selective medium without tryptophan, leucine, and histidine selects for the interaction between bait and prey.

### Cloning β-glucosidases and performing enzymatic assays

The β-glucosidases enzymatic assays were performed as described in^45^. Briefly, cDNAs for *bGLU23*, *bGLU25* were amplified from *A. thaliana* shoot cDNA library using specific primers detailed in Table S2. bGLU25^G420E^ sequence was synthesized (Genscript). The DNA products were ligated into the pBluescript SK(+) vector and transformed into *E. coli*. After plasmid extraction, sequencing confirmed the presence of the insertion sequences. The inserts were excised, gel-purified, and then cloned into the pET28 expression vector. Recombinant proteins were purified through nickel affinity chromatography. For β-glucosidase activity assays, we used the Millipore Sigma Kit (MAK129) and followed the recommendation.

### Subcellular Fractionation and Immunoblot Analysis in Arabidopsis and Tobacco

To separate cytosolic, ER-enriched, or nuclear fractions, Arabidopsis plants were homogenized and fractionated using either the Minute™ Plant Microsomal Membrane Extraction Kit (Invent Biotechnologies, MM-018) or the Minute™ Plant Cytosolic and Nuclear Protein Isolation Kit (Invent Biotechnologies, PF-045), following the manufacturer’s instructions. Total protein extraction was performed by grinding 100 mg of tobacco leaves and Arabidopsis shoots in liquid nitrogen, followed by suspension in extraction buffer (50 mM Tris–HCl pH 7.5, 150 mM NaCl, 10% glycerol, 5 mM dithiothreitol, 2 mM Na₂MoO₄, 2.5 mM NaF, 1.5 mM activated Na₃VO₄, 1 mM PMSF, 1% IGEPAL, and 1X cOmplete™ Protease Inhibitor Cocktail (Roche)). Soluble proteins were obtained by centrifuging cell debris three times at 12,000 rpm for 10 min at 4°C. Proteins were denatured with 6× Laemmli buffer at 95°C for 5 min, and 25 µg of protein was loaded into each well of 10% or 8–16% SDS-PAGE gels and transferred to PVDF membranes (Bio-Rad, 1620174). Membranes were blocked with 5% skim milk for 1 h at room temperature. For immunoblotting, a GFP antibody (Miltenyi Biotec, 130-091-833) diluted 1:3000 was incubated overnight at 4°C. Secondary anti-rabbit IgG (Sigma-Aldrich, A0545) diluted 1:10,000 was incubated for 2 h at room temperature. ER, cytosolic, and nuclear markers were assessed using antibodies for Lumenal-binding protein (BiP; Agrisera, AS09481), UDP-glucose pyrophosphorylase (UGPase; Agrisera, AS05086), α-Tubulin (Sigma-Aldrich, T6199), and Histone H3 (Agrisera, AS10710), all diluted 1:2000 and incubated overnight at 4°C. Secondary anti-rabbit IgG or anti-mouse IgG (Sigma-Aldrich, A9044, 1:10,000) was used. Immunoblots were detected using Clarity Western ECL Substrate (Bio-Rad, 1705061) and visualized by a Bio-Rad ChemiDoc system.

### Protein interaction analysis

*Agrobacterium*-mediated transient expression was performed using the Agrobacterium GV3101 strain as described. Briefly, overnight-grown *Agrobacterium* culture was resuspended in induction medium (10 mM MES-KOH, pH 5.7, 10 mM MgCl_­_, and 100 μM acetosyringone) to OD_­_= 0.2 and incubated for 2 hours at room temperature, before infiltrating *N. benthamiana* leaves. *Agrobacterium* strain carrying the 35Spro:p19 construct was co-infiltrated to enhance the level of protein expression. Transiently expressed proteins were analyzed 2 days after infiltration with a Nikon A1R confocal laser scanning microscope. Co-IP assays were performed as described^46^. The ssGFP-bGLU25 sequence was synthesized and cloned into the pUC57 vector via Genscript. Using LR clonase, bGLU25 was transferred into the pMDC32 vector and introduced into Agrobacterium tumefaciens strain GV3101. For ΔbGLU25-GFP, the coding sequence excluding 45 nucleotides was amplified with specific primers (Table S3), and cloned into pDONR223. This was transferred into pMDC83 under the 35S promoter and introduced into *A. tumefaciens*. The full-length AtJAC1 coding sequence was amplified with specific primers (Table S3), cloned into pDONR223, transferred to the pEarlyGate 101 vector, and introduced into A. tumefaciens. ΔbGLU25-GFP or ssGFPbGLU25 was coexpressed with AtJACI-HA in *N. benthamiana* leaves. After 2 to 3 days, plant material was harvested and ground in native protein extraction buffer, filtered, and centrifuged. The supernatant was incubated with anti-GFP antibody coupled to protein A/G Sepharose beads (Millipore), the beads were washed four times with wash buffer, and bound proteins were eluted with elution buffer containing 2% (w/v) SDS for immunoblotting with HA antibody.

### Confocal laser scanning microscopy and image analysis

Confocal image acquisition was performed using an inverted laser scanner confocal microscope Nikon A1Rsi on *Arabidopsis* and *N. benthamiana* leaves. The fluorescent proteins used in this study were EGFP, RFP and (m)Venus (Clontech, http://www.clontech.com/). BiFC assays were performed as described in^46^. Briefly, constructs for the expression of bGLU25 or ΔbGLU25-mVenus^N^ or-mVenus^C^, GRP7-mVenus^N^, and AtJAC1-mVenus^c^ were generated and infiltrated into *N. benthamiana* leaves YFP fluorescence (suitable for mVenus: excitation filter BP 500/20, dichromatic mirror 515, BP suppression filter 535/30) was visualized. Tobacco leaves were infiltrated and transiently transformed combining the constructs to be tested for association using a confocal microscope (Nikon A1Rsi CLSM) 40 to 48 h after infiltration. Imaging results presented in this work are representative of at least three independent experiments.

### Gene expression analysis by quantitative RT-PCR

Gene expression analysis involved extracting total RNA from 100 mg of frozen *Arabidopsis* shoot samples using the Plant RNeasy extraction kit from Qiagen (model number). RNA concentrations were determined with a NanoDrop spectrophotometer (Thermo Scientific, model number). For cDNA synthesis, 2 μg of RNA was utilized. Real-time PCR was performed using a LightCycler 480 PCR System (Roche) with a 20 μL reaction mixture containing 10 μL of SYBR Green I master mix, 0.3 μmol of each primer (Table S3), and 5 μL of a 1:25 dilution of cDNA. The PCR protocol included an initial denaturation at 95°C for 5 minutes, followed by 42 cycles of 95°C for 10 seconds, 60°C for 10 seconds, and 72°C for 25 seconds, finishing with a final extension at 72°C for 5 minutes. The expression levels of *bGLU25*, *SCP50*, and *FLC* mRNA were quantified and normalized against *UBQ10* mRNA, with data presented as relative expression levels compared to wild-type and mutant samples as described in^47^. The same method was used to quantify OsbGLU33 expression in rice, using *UBQ10* mRNA as a reference for relative expression analysis.

### Statistical analysis

Statistical analysis of quantitative data was performed using GraphPad Prism 9 software for macOS. A p-value < 0.05 was considered statistically significant for all t-test analyses. Student’s two tailed t test was used for statistical analysis, assuming equal variance, and data with P value 0.05 were considered significant. Pearson’s correlation analyses were used to estimate signal overlap, and one-way ANOVA was used for statistical analysis.

## ACKNOWLEDGMENTS

Funding: This work was funded in part by the National Science Foundation (NSF, USA), Div Of Molecular and Cellular Bioscience (MCB-233456), Michigan State University (USA) to H.R. and F.B., and the Plant Resilience Institute to H.R; Lahore University of Management Sciences, Pakistan to Z.S., and US National Science Foundation grants (IOS-2312181, IOS-2406533, IOS-1546838, DBI-2213983) and US Department of Energy, Office of Science, Office of Biological and Environmental Research, Genomic Science Program grants (DE-SC0018277, DE-SC0020366, DE-SC0023160, and DE-SC0021286) to S.Y.R. The funders did not have any role in study design, data collection, and analysis, decision to publish, or preparation of the manuscript.

## AUTHOR CONTRIBUTIONS

H.R. and S.Y.R. conceived the project. Experiments designed by H.R. H.C., A.N, Z.S., and N.B. performed the genome-wide association mapping on flowering time under phosphorus limitation. N.B., I.C., A.N., L.Z., and H.C. performed the qRT-PCR analyses, generated plasmid constructs, the homozygote mutants, and the complemented mutant lines in Arabidopsis. H.R. S.Y.R., F.B., and Z.S. wrote the manuscript. Competing interest: The authors declare no competing interests. Data and materials availability: Data supporting the findings of this work are available within the paper along with its Supplementary Information and Source Data files. The datasets and plant materials generated and analyzed during this study are available from the corresponding author H.R. upon request.

## SUPPLEMENTARY FIGURES

**Supplementary Figure 1.**
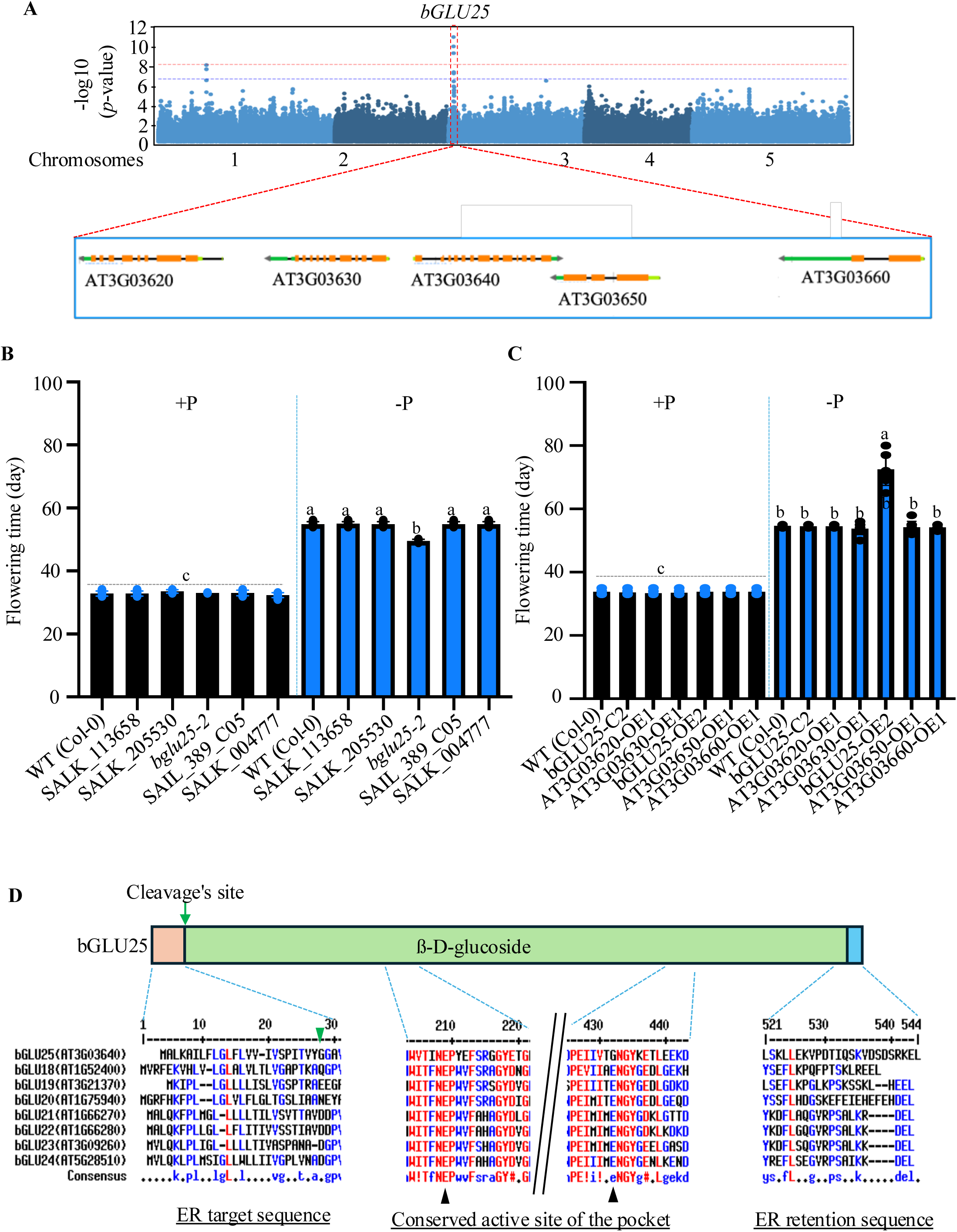
bGLU25: Adapting Flowering in *A. thaliana* to Phosphorus Availability. **(A)** Gene loci located within 20 Kb of the significant SNP on chromosome 3 (Chr3:947345), which is highlighted in a red rectangle. **(B)** Flowering time data for Arabidopsis Col-0 (WT), *bglu25* knock-out mutant (*bglu25-2*), T-DNA insertion mutants SALK_113658 (AT3G03620), SALK_205530 (AT3G03630), SAIL_389_C05 (AT3G03650), and SALK_004777 (AT3G03660), grown under +P and -P conditions. **(C)** Flowering time data for Arabidopsis Col-0 (WT), complemented *bglu25-1* (bGLU25-C2) the overexpressor lines of AT3G03620, AT3G03630, bGLU25 (AT3G03640, bGLU25-OE2), AT3G03650, and AT3G03660 driven by the 35S promoter, grown under +P and -P conditions. **(D)** bGLU25 contains an endoplasmic reticulum (ER) targeting sequence (1-23) and ER retention sequence, represented by a KDEL tail at the C-terminus. Comparison of the bGLU25 protein sequence with other β-glucosidases from bGLU18 to bGLU24 in Arabidopsis revealed that bGLU25 lacks the functionally important second catalytic residue (420E) but the first active site (203E) was conserved. The alignment used through http://multalin.toulouse.inra.fr/multalin/help.html#Output_formats site. The three colors (red, blue and black) for the sequence residues present the high consensus, low consensus, and neutral. A green triangle indicates cleavge site after ER target signal, and black triangles mark important conserved regions (1’st proton donor, and 2^nd^ nucleophile sites) in the catalytic site of arabidopsis bGLU in the same group. E: Glutamic Acid.

**Supplementary Figure 2.**
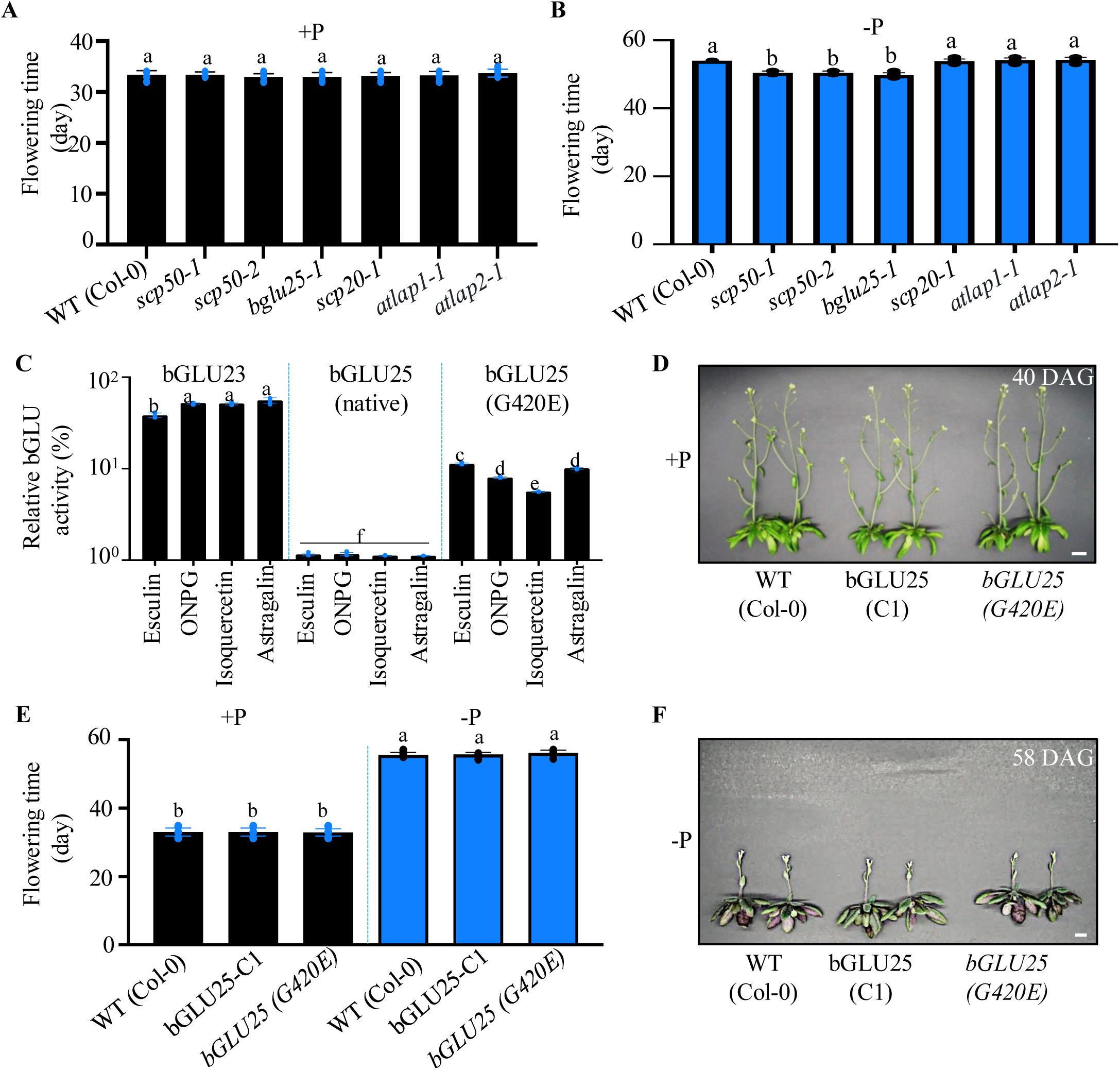
Cytosolic Translocation of bGLU25 Under Phosphorus Limitation is Mediated by SCP50, Crucial for Delaying Flowering Time. **(A)** and (**B**) Flowering times of wild-type (WT) plants (Col-0) and several mutants: *SERINE CARBOXYPEPTIDASE 20* (*scp20-1*), *SERINE CARBOXYPEPTIDASE 50* (*scp50-1*), *scp5-2, LEUCINE AMINOPEPTIDASE 1* (*atlap1-1*), *LEUCINE AMINOPEPTIDASE 2* (*atlap2-1*), and *bGLU25 (bglu25-1)* under the +P (**A**) and -P (**B**) conditions. The letters indicate significant differences in flowering times (p < 0.05), determined using one-way ANOVA followed by a Duncan post hoc test. **(B)** Enzymatic activity of β-glucosidases: active bGLU23, inactive bGLU25, and mutated bGLU25 (G420E). Reactions used esculin, ONPG, isoquercetin, and astragalin, repeated three times. Results are shown as Relative bGLU activity (%). G: Glycine, E: Glutamic Acid. **(C)** and (**F**) Representative images of wild-type (WT) plants (Col-0) and *bglu25* mutant plants (bGLU25-C1) that were complemented with either native bGLU25 or mutated bGLU25 (G420E). The plants grown under +P conditions (**D**) were photographed at 40 days (scale bar = 1 cm), while those grown under -P conditions (**F**) were photographed at 58 days (scale bar = 1 cm). **(D)** Flowering time data of WT (Col-0) and *bglu25* mutant (bGLU25-C1) plants complemented with either native bGLU25 or mutated bGLU25 (G420E), grown under +P (**D**) and -P (**F**) conditions. Results are presented as means ± SD (n = 18), with letters indicating significant differences (p < 0.05).

**Supplementary Figure 3.**
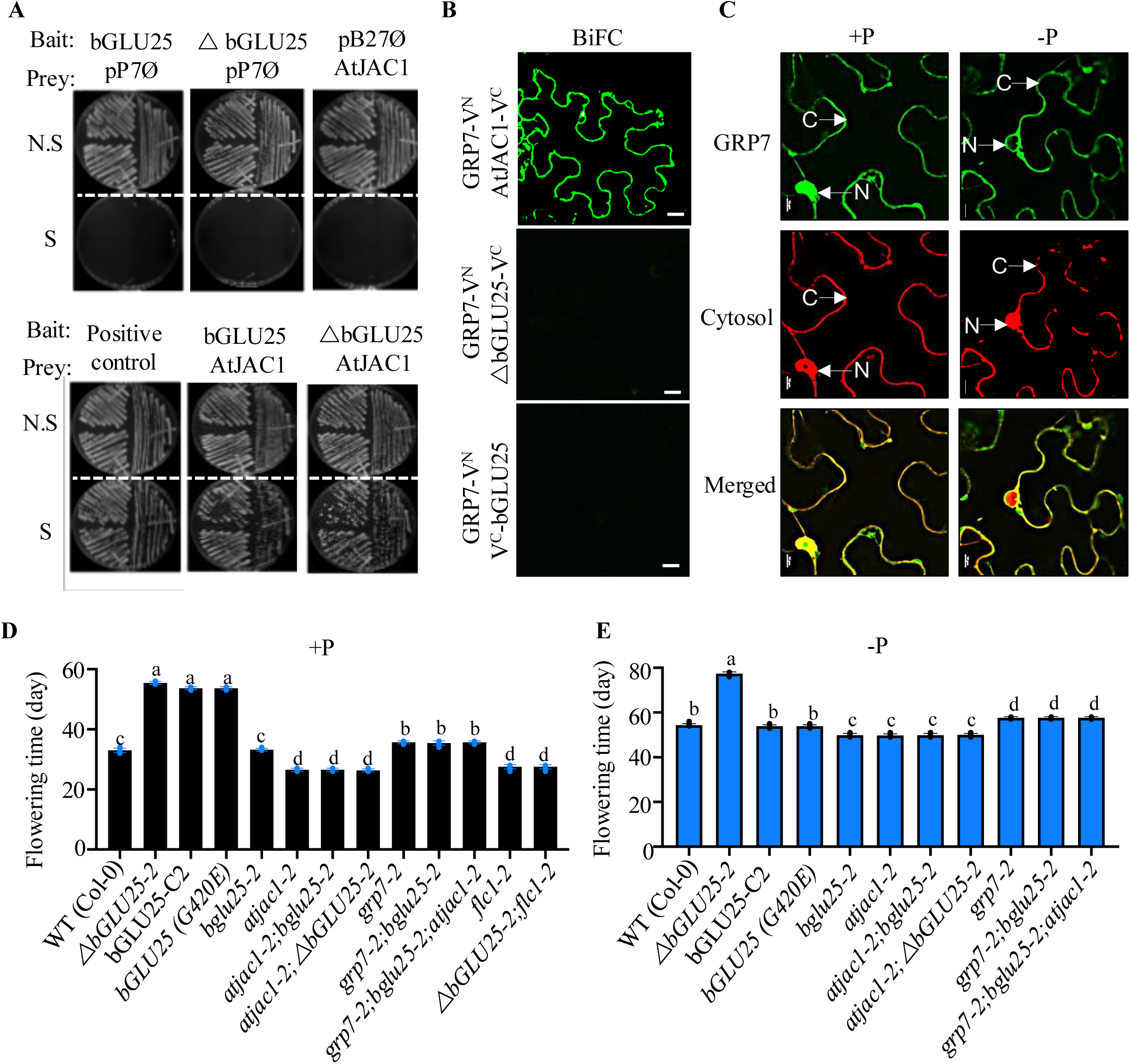
The bGLU-AtJAC1-GRP7-FLC Pathway Regulates Flowering Time in *Arabidopsis*. **(A)** Yeast Two-Hybrid (Y2H) analysis of an *A. thaliana* shoot library identified JACALIN-LECTIN LIKE1 (AtJAC1) as an interactor of both the full-length bGLU25 and ΔbGLU25. This is shown by growth on selective media (S, DO-3) lacking tryptophan, leucine, and histidine. Positive controls included SMAD (Suppressor of Mothers against Decapentaplegic) and SMURF (SMad Ubiquitination Regulatory Factors) from the human TGF-β (Transforming Growth Factor-β) /Smad pathway, while negative controls were bGLU25, ΔbGLU25 or AtJAC1 alone. **(B)** The BiFC assay tested GRP7-mVenus^n^ interaction with bGLU25-mVenus^c^ or ΔbGLU25-mVenus^c^ in *N. benthamiana*, using GRP7 and AtJAC1 interaction as positive controls (scale bar = 10 µm). **(C)** Transient expression of GRP7 in *N. benthamiana* leaves alongside a cytosolic marker, cRFP, under varying P conditions (+P and -P). Under -P conditions, GRP7 is mainly found in the cytosol (scale bar = 10 μm). C, cytosol. N, Nucleus. **(D)** Flowering time data for various genotypes grown under +P conditions, including wild type (Col-0), *ΔbGLU25-2*, a complemented *bglu25-1* line with *bGLU25* (bGLU25-C2), and others. Results are presented as means ± SD (n = 18), with letters indicating significant differences (p < 0.05). **(E)** Flowering time data for the same genotypes grown under -P conditions, with means ± SD (n = 18) and letters indicate significant differences (p < 0.05).

**Supplementary Figure 4.**
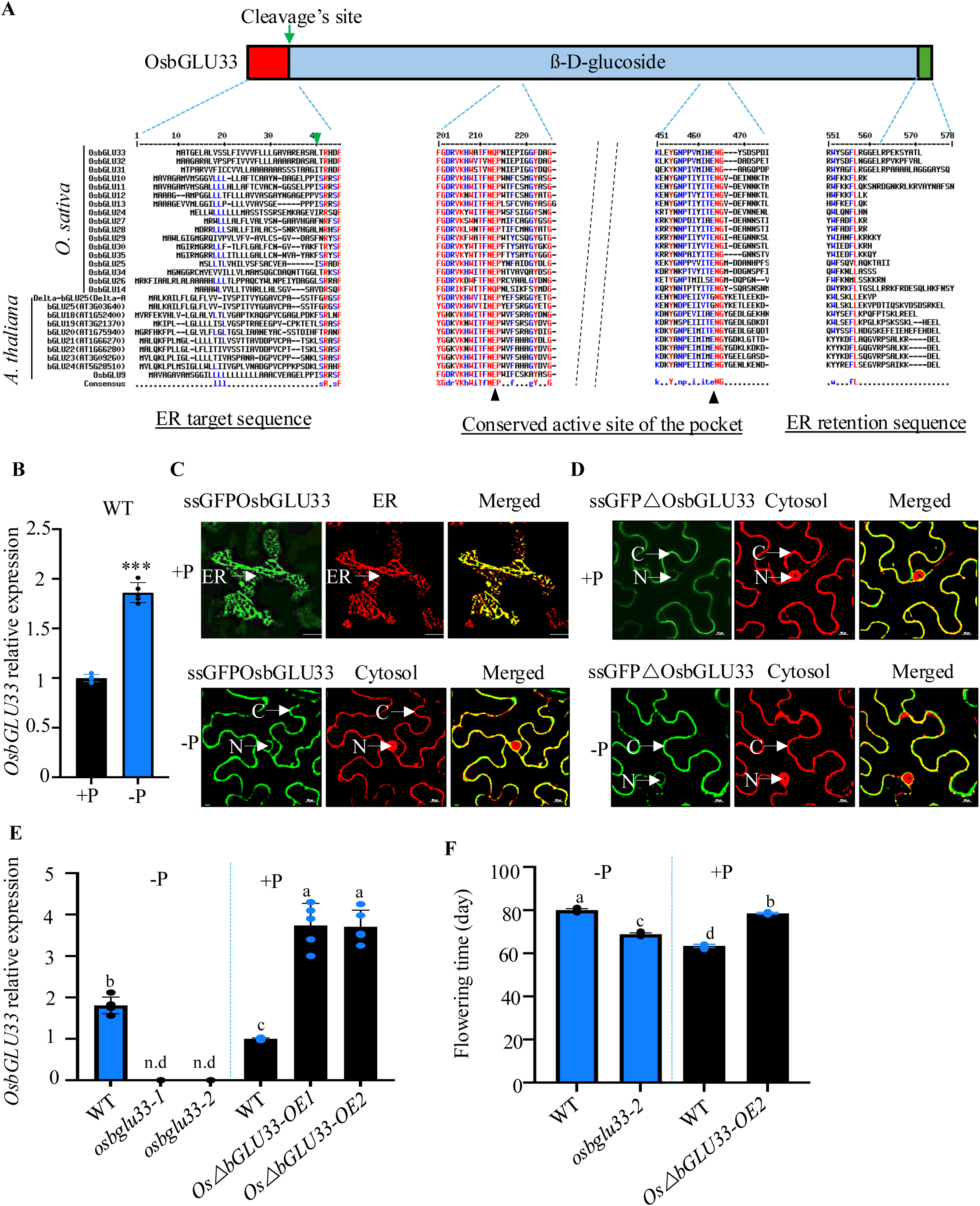
The bGLU-Dependent Regulation of Flowering Time is Shared in Rice. **(A)** OsbGLU33 contains an ER targeting sequence (1-30) and ER retention sequence, represented by a KDEL tail at the C-terminus. Comparison of the OsbGLU33 protein sequence with other rice and Arabidopsis β-glucosidases through http://multalin.toulouse.inra.fr/multalin/help.html#Output_formats site. **(B)** Relative mRNA abundance of *OsbGLU33* in 65-day-old *O. sativa* shoots was quantified by qRT-PCR, normalized to *Ubiquitin10*. Means (±95% confidence interval). 9 plants from 3 independent experiments were analyzed. Student’s t test, \*\*\**p* < 0.001. **(C)** Confocal microscopy images of *N. benthamiana* leaves from plants grown under +P and -P conditions show the GFP signal from ssGFPbGLU33 (green fluorescence, left), ER marker HDEL-RFP (red fluorescence, middle) or the cytosol marker (red fluorescence, middle), and an overlay of both images (right). Scale bar: 10 μm. ER, Endoplasmic Reticulum. C, cytosol. N, Nucleus. **(D)** Confocal microscopy images of *N. benthamiana* leaves from plants grown under +P and -P conditions show the GFP signal from ssGFPΔbGLU33 (green fluorescence, left), the cytosol marker (red fluorescence, middle), and an overlay of both images (right). Scale bar: 10 μm. C, cytosol. N, Nucleus. **(E)** Relative mRNA abundance of *OsbGLU33* in the shoots of 65-day-old rice plants grown under +P and -P conditions, quantified by qRT-PCR and normalized to *Ubiquitin 10*. Significant differences are indicated. **(F)** Flowering time of WT, *osbglu33* knock-out mutant (*osbglu33-2*), and Os*ΔbGLU33* overexpressor (*OsΔbGLU33OE-2*) plants grown under +P and -P conditions, with significance indicated.

